# Evaluating Named-Entity Recognition approaches in plant molecular biology

**DOI:** 10.1101/360966

**Authors:** Huy Do, Khoat Than, Pierre Larmande

## Abstract

Text mining research is becoming an important topic in biology with the aim to extract biological entities from scientific papers in order to extend the biological knowledge. However, few thorough studies on text mining and applications are developed for plant molecular biology data, especially rice, thus resulting a lack of datasets available to train models able to detect entities such as genes, proteins and phenotypic traits. Since there is rare benchmarks for rice, we have to face various difficulties in exploiting advanced machine learning methods for accurate analysis of rice bibliography. In this article, we developed a new training datasets (Oryzabase) as the benchmark. Then, we evaluated the performance of several current approaches to find a methodology with the best results and assigned it as the state of the art method for our own technique in the future. We applied Name Entities Recognition (NER) tagger, which is built from a Long Short Term Memory (LSTM) model, and combined with Conditional Random Fields (CRFs) to extract information of rice genes and proteins. We analyzed the performance of LSTM-CRF when applying to the Oryzabase dataset and improved the results up to 86% in *F*_1_. We found that on average, the result from LSTM-CRF is more exploitable with the new benchmark.

## 1 Introduction

### 1.1 Problem Statement

The last few decades have witnessed the massive explosion of information in life science. However, an important proportion of information relevant to this field is not available from databases but is instead present in unstructured scientific documents, such as journal articles and reports. Agronomy is an overarching field, that comprises of diverse domains such as genetics, plant molecular biology, ecology and soil science. Despite the advancement in information technologies, scientific communication in agronomy is still largely based on text because it is the common way to report scientific advancements. To effectively develop applications to improve crop production through sustainable methods, it is important to overlay research findings from these fields as they are highly interconnected. However, the collection of content is growing continuously and the information are currently available as unstructured text. Using these resources more efficiently and taking advantage of associated cross-disciplinary research opportunities poses a major challenge to both domain scientists and information technologists.

Several text mining methods and tools have been developed to solve the problem of named entity recognition by using different approaches [14]. However, to handle with identifying biological entities issues, there are four possible and common approaches [2].

– The most basic and traditional methodology is rule-based approach. The technique identifies entities by a group of written rules, which are manually done by the domain scientists with linguistic knowledge. As a result, it is time-consuming and easily error-prone.
– The dictionary-based is common method to handle NLP problems. The model matches the candidate entities with a dictionary that contains all the known entities to detect whether the candidate belongs to a defined category or not. However, if there exists new entities, for instances, from new discovery and not in the available dictionary, the system is not able to recognize them, which reduces the efficiency of the model.
– The third approach is based on machine learning, which uses a statistical classifier to extract the features (prefixes, suffixes, number of capital letters, etc.) that are able to recognize the entities. Several familiar algorithms have been proposed such as Naive Bayes, Conditional Random Fields (CRFs) [10], and so on. However, this method need an important corpus annotated manually to train and test efficiently.
– Hybrid approach is proposed as the combination of two or more of the previous techniques to take advantage of the strength and reduce the weak points among them.

Identifying biological entities is not trivial. Despite the fact that there exists many available approaches to handle this problem in general and in the Biomedical domain, few thorough studies have been implemented for plants, especially rice. Moreover, we found rare benchmarks available for plant species and none for rice. Thus, we faced several difficulties exploiting advanced machine learning methods for accurate analysis of rice.

### 1.2 Objective

In the large scale, we are currently building an RDF knowledge base, Agronomic Linked Data (AgroLD – www.agrold.org). The knowledge base is designed to integrate data from various publicly plant centric databases such as Gramene [13] and Oryzabase [16] to name a few. The aim of AgroLD project is to provide an integrated portal for bioinformatics and domain experts to exploit the homogenized data model towards filling the knowledge gaps. In this landscape, we aim to extract relevant information from the scientific literature in order to enrich the content of integrated datasets.

Due to the scope of the article, we exploited information from Oryzabase database as our benchmark and focused on researching and evaluating the performance of the current approaches and assigning the method with the best accuracy and efficiency as the baseline for our own technique in the future. Thus, we identify two relevant approaches: Long Short Term Memory - Conditional Random Fields (LSTM-CRF) and the hybrid method between dictionary lookup and machine learning algorithms, has been chosen for further analysis due to their competency and efficiency compared to others.

## 2 Materials and methods

### 2.1 NER-tagger with LSTM-CRF

Long Short Term Memory - Conditional Random Field (or LSTM-CRF) is a generic method based on deep learning and statistical word embeddings [7]. To deeply understand the whole architecture, it is necessary to understand the concept of LSTM and CRF first.

#### LSTM

When Recurrent Neural Network (RNN) are applied in practice for sequence prediction, they tend to fail to predict input that are depended on long previous sequences. This problem called “The long-term dependencies” explored by Hochreiter [8] and Bengio *et al*. [3] Long Short Term Memory, a special kind of RNN, was introduced to avoid the long-term dependency problem of RNN by Hochreiter & Schmidhuber (1997) [9]. With the addition of three sigmoid layers, it becomes more complex than a traditional RNN but it proves that it works effectively to solve the problems of long-term dependencies. The sigmoid layers with output between [0,1] decide which information will be forgotten while the tanh layer use that output to create a vector of candidates during the training process [9]. Taking a sequence of vectors (*x*_1_, *x*_2_, …, *x_t_*) as an input, LSTM returns the output in the form of another sequence (*h*_1_, *h*_2_, …, *h_t_*) of equal length to represent the information of the input. These equations belows describe the implementation of LSTM:

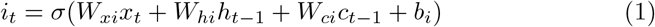

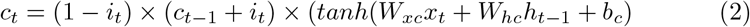

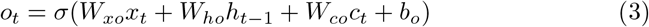

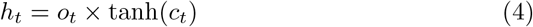

where *σ* is the element-wise sigmoid function, and *×* is the element-wise product.

The workflow follows the combination of LSTM pair (forward and backward LSTM) called a bidirectional LSTM (bi-LSTM)[6]. After receiving the input (*x*_1_, *x*_2_, …, *x_t_*) where each element is represented as a d-dimensional vector, the forward LSTM layer computes the representation 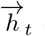 of the left context of the sentence at every word t while the backward LSTM reads the same input sequence in reverse to obtain the representation 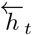 of the right context using different parameters. In the next step, the model concatenates its left and right context representations, 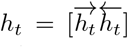 to achieve the representation of a word, which is useful for various tagging applications.

#### CRF

Conditional random fields (CRFs), introduced by Lafferty *et al*. [10], is a statistic modeling method for labeling and segmenting structured data, such as sequences, trees and lattices. The general idea of this model [12] is to make independent tagging decisions for each output *y_t_* using the features *h_t_*. However, for NER tagger, using traditional CRFs with independent classification decisions is insufficient and impossible with numerous constraint because of the strong dependencies cross the output labels.

In this article, we focus on the jointly CRFs model. For an input *X* = (*x*_1_, *x*_2_, …, *x_t_*), the matrix of scores output P is obtained from the bi-directional LSTM network with *n* x *k* in size, where k is the number of tags and *P_i,j_* regarding the score of *j^th^* tag of *i^th^* word in a sentence [11]. Meanwhile, for the sequence of predictions *Y* = (*y*_1_, *y*_2_, …, *y_t_*), the score is:

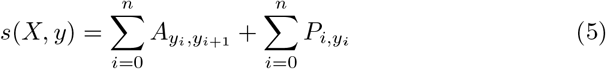

where A represents a [k + 2] matrix of transition scores from the tag *y_i_* to tag *y_j_*.

The probability for the sequence y then is de_ned by a softmax function over all possible tag sequences as:

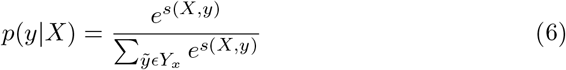

To encourage the model to produce a valid sequence of output labels, the log-probability of the correct tag sequence is maximized in training process as:

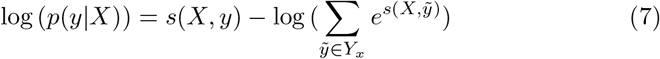

where *Y_X_* indicates all possible tag sequences for a sentence X.

After that, we predict the output sequence in decoding step following the equation:

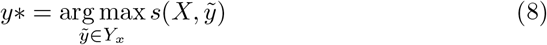

#### LSTM-CRF

The architecture of LSTM-CRF is illustrated in Fig.1 [7]. The whole system includes three main layers: the embedding layer as input, bidirectional LSTM layers, and a CRF layer as output. Given the raw sentence made of the sequence of words *w*_1_; *w*_2_; …; *w_n_* as the input, the embedding layer produces an embedding vector *x*_1_; *x*_2_; … ; *x_n_* for each word. Every embedding vector regarding distinct word is a concatenation of two components: wordand a character-level embedding. We retrieved the word-level embedding from a lookup table of word embedding, meanwhile, we applied a bi-directional LSTM to the character sequence of each word and then concatenate both directions to achieve the character-level embedding. That means the resulting sequence of embeddings *x*_1_; *x*_2_; … ; *x_n_* is fed into bi-directional LSTM-layer to produce a refined representation of the input and we take as the input to a final CRF layer. The final output from this layer is obtained by applying the classical Viterbi algorithm. We trained the whole network by back-propagation.

**Fig. 1.**
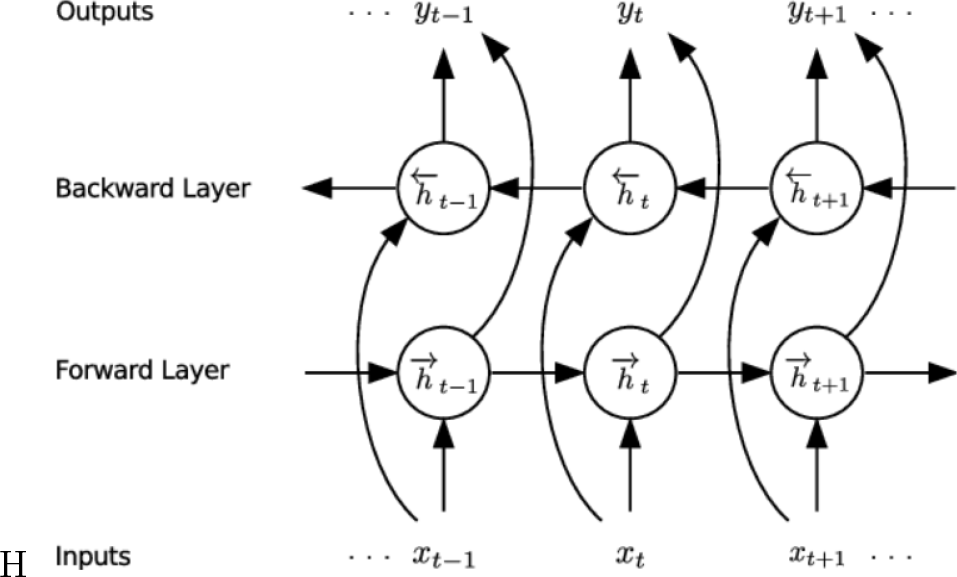
LSTM model [5].

**Fig. 2.**
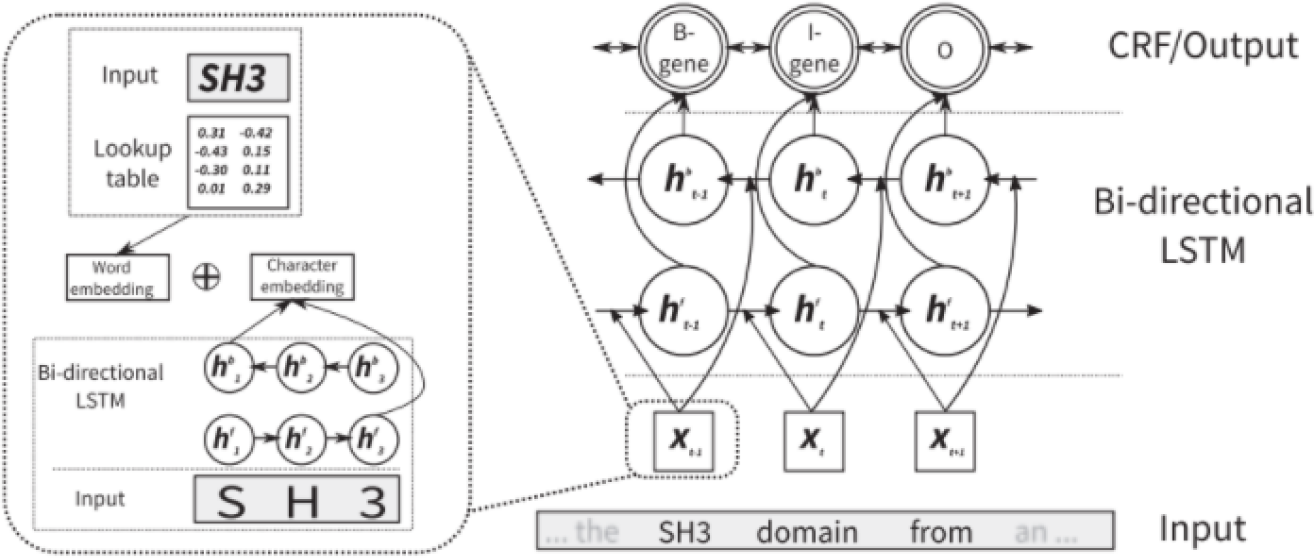
LSTM-CRF model[7].

### 2.2 NER-hybrid approach

This approach is based on a dictionary lookup entity recognizer combined with machine learning classifiers. The OGER entity recognizer is used to annotate the objects in some selected domain ontologies. Next, the Distiller framework is applied to extract this information as a feature for a machine learning algorithm to select relevant entities. For this step, we implemented two different machine learning algorithms: Conditional random fields and Neural networks (NNs).

#### OGER

The Onto Gene group has developed an approach for biomedical entity recognition based on dictionary lookup and flexible matching [2]. Their method has been applied in several competitive evaluation of biological text mining technologies, and often archived good results.

OGER performs a flexible interface for dictionary based annotation. It provides the annotated terms along with the corresponding identifiers either in a simple tab-separated text file. For the given input which is Oryzabase references included over 10000 abstracts and titles of scientific articles, we implemented OGER as a permanent web service which is accessible through an API. Before implementation, input data was extracted into several files, with the same format to make the evaluation process be more easily. OGER architecture is configured to have a strategy towards to a greater false positives (FP) which leads to a lower precision value, but less false negatives (FN) which can get an high recall. Because the aim of the project focus on the information of plant genes and proteins and their relationship, we chose the gene dictionary of Gene Ontology Consortium [4] and the protein dictionary of [15] came from back-end software of Bio Term Hub, which is a meta-resource for biological resources. In OGER, the entities of the term dictionary were then preprocessed in the same way as the documents with respect to tokenization, stemming, and case sensitivity, as described below. Next, the input documents will be compared to the dictionary with an exact-match strategy.

#### Distiller

The Distiller framework is an open-source project which is developed to build a flexible, extensible system for natural language processing fields [2]. The main process of Distiller based on the performance of Automatic Key-phrase Extraction (AKE) to extract information from text. AKE seems to be different from NER, as while the former is focus on finding the small set of the most relevant information in a document, and then find all the information of selected types. Besides, AKE can be performed both as an unsupervised and supervised algorithms, and Distiller actually derives from an unsupervised approach.

About the architecture, Distiller is organized in series of single-knowledge oriented modules where each module is designed to perform a single task efficiently [1] such that part-of-speech (POS) tagging, statistic analysis, and so on. It performs with the ability to implement various pipelines for different tasks. All modules are required to share the entity extracted results on a ≪blackboard≫, thus a module can share the knowledge with other ones. Implementing KE tasks with Distiller is reduced to specifying pipeline with annotators [1]. A task is normally divided into steps: text pre-processing, candidate key-phrase generating, and candidate raking. The pipeline process of Distiller is described in figure 4.

In this case, we performed AKE methods using supervised machine learning algorithms. Normally, the first step generates the key-phrases, which use POS tagger to select patterns. Then, we assigned features to keyphrase, using statistical, linguistic, or semantic knowledge. Finally, a machine learning algorithm is trained and then evaluated over a set of documents associated with ground truth.

However, in our case, we replaced the key-phrase generating step by OGER’s output.

#### Models and Features

In our project, we proposed two machine learning algorithms: neural network and conditional random fields which are considered to perform the exploitable results [2]. The performance of Distiller is different between two methods. In case of CRF, it uses the annotated output of OGER as a feature, and considers any token in the text as an entity to predict. In contrast, Distiller only focus on filtering OGER’s output, and process to classify for each entity phrase. The workflow for the hybrid approach will be implement as follows:

1. Based on traditional dictionary lookup, tag object from text data using OGER from given input.
2. Integrate output of OGER with Distiller framework to assign new features, prepare them to be processed by a machine learning system.
3. Build a machine learning system including a simple neural network and a CRF, select the relevant entities in the document using the information generated in the previous steps.

For CRF, we considered each word as a token and tried to add some features. To do some experiments, we trained several models with different kinds of features. Finally, we selected a number of features having an effect to the performance of CRF models. The list of CRF features is showed in the table 2.

**Table 1.** The details of CRF features.

| Conditional Random Fields |  |
| --- | --- |
| Input | Single tokens |
| Features |  |
| Candidate POS-tagger | Yes |
| Candidate tag (IOB format) | Yes |
| Candidate contains numbers | Label yes/no |
| Candidate ends with numbers | Label yes/no |
| Candidate pre-selected by OGER | Label yes/no |
| Neighbor tokens | Yes |
| Neighbor token POS-taggers | Yes |
| Total features for each tokens | 23 |

**Table 2.** The details of NN features.

| Neural Network |  |
| --- | --- |
| Input | n-grams selected by OGER |
| Features |  |
| Candidate tag (IOB format) | Yes |
| Distiller features | Yes |
| Candidate contains numbers | Count |
| Candidate ends with numbers | Label yes/no |
| Total features | 13 |

For NN, we considered input as n-grams that selected by OGER. The features are generated from Distiller and some manual generated features. The list of CRF features is showed in the table 3.

**Table 3.** The details of main dataset.

| Dataset | Text genre | Text type | Entity Type |
| --- | --- | --- | --- |
| Oryzabase | Scientific article | Abstract and title | Genes |

**Table 3.** The details of main dataset
| Number of articles | Number of sentences | Number of words |
| --- | --- | --- |
| 10400 | 75096 | 2697726 |

### 2.3 Dataset

In this project, we built and implemented some approaches on datasets collected from Oryzabase^4^, an integrated scientific database for Oryza sativa (or rice) species published online since 2, 000 [16]. The lasted version of Oryzabase contains 21, 739 of rice genes, collected from 44, 837 distinct scientific articles. The PubMed database is used as resource to collect the raw text which is preprocessed later to form the input data. However, a number of scientific articles were not found on the PubMed database due to the published resource and some historical issues. After filtering, over 10, 000 articles were processed. The detailed of raw data is shown in table 3. Next, concentrating on the entities of rice genomes, we took the Oryzabase gene list as the ground truth to built our dataset by keyword matching term respectively.

## 3 Implementation

### 3.1 Text pre-processing

Due to the size of data, we only focused on the title and abstract of each article, which might contain a lot of information about genes and their relationship such as gene name, gene symbols or gene symbol synonyms, etc. The usage of whole articles may lead to the noise of data after process and the waste of time memory during the training phase. The original format of data from PubMed were formed into separated tabs, which include the pubmed ID, title, released year, journal and content of abstract.

The first step is to convert the raw data into a uniform format. To tokenize data, each word is considered as a token, given in the following lines; one token per line, and included 3 tabs: the first one being the word itself, the part-of-speech tags (POS tag) and the last one being the entity type. Each token is on a separate line, ends with dots and has an empty line after each sentence. In addition, an empty line indicates the end of a document, and the next document starts after this empty line.

To do that, raw data is split into sentences, and from sentences into tokens. Next, we matched words with the Oryzabase list gene dictionary by keys and applied the IOB (Inside-Outside-Beginning) format to determine the token be entity or not. To make the tag become more meaningful for CRF, POS tag is used on each token by a library for natural language processing. During the implementation of two methods, this pre-processed data is used to build and train the model with some additional features based on different purposes. To avoid some unexpected errors during the training part, data is converted into lower case; and some “non-alphabet” character is ignored.

### 3.2 NER-tagger

Before going to training process, we need to define some properties. First, the LSTM (with CRF) will work at word embedding level which include an entity tag to determine itself and the input data going through model is embedding sequences. The LSTM (with CRF) used for NER built based on the model of Lample *et al* [11]. To start the training process, we have done several experiments to find the best parameters for this model. Some results were very exploitable at a specific value of learning rate and dropout. Besides, we tried an LSTM model with and without the CRF to evaluate the performances on general results.

### 3.3 Hybrid Methods

For the hybird approach, OGER, a dictionary tool supported by Gene Ontology and Protein Ontology dictionary, used to annotate all objects in the dataset. The output was formatted into tables which contains full information of genes. Next, some first evaluations were applied to evaluate the performance of OGER on the whole dataset. The purpose of this evaluation was to see how OGER effectively work on this dataset and based on that to improve the results.

For CRF and NN, we also started with POS-tag and candidate tag prepared in the pre-processsing part and combined with new features that are generated and listed above.

### 3.4 Evaluation parameters

We realized that the Oryzabase dataset is quite big, thus we decided to split it into 3 subsets with different ratio: 75% for training set, 15% for testing set, and the last 10% of dataset for validation set which can help the model avoid over-fitting issue (Fig.3).

**Fig. 3.**
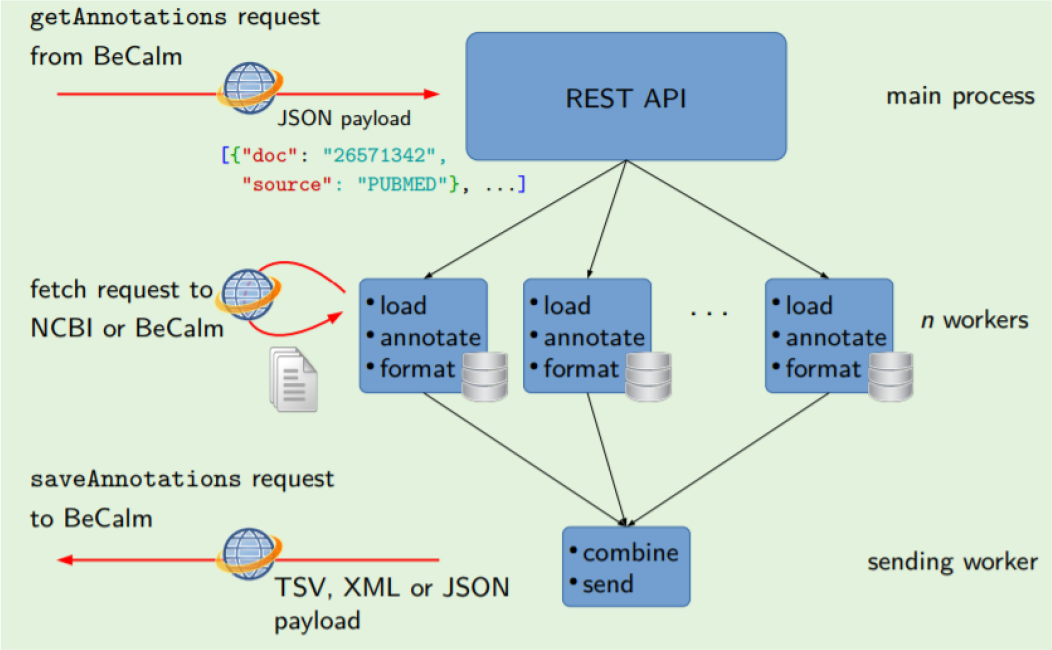
the structure of OGER.

**Fig. 4.**
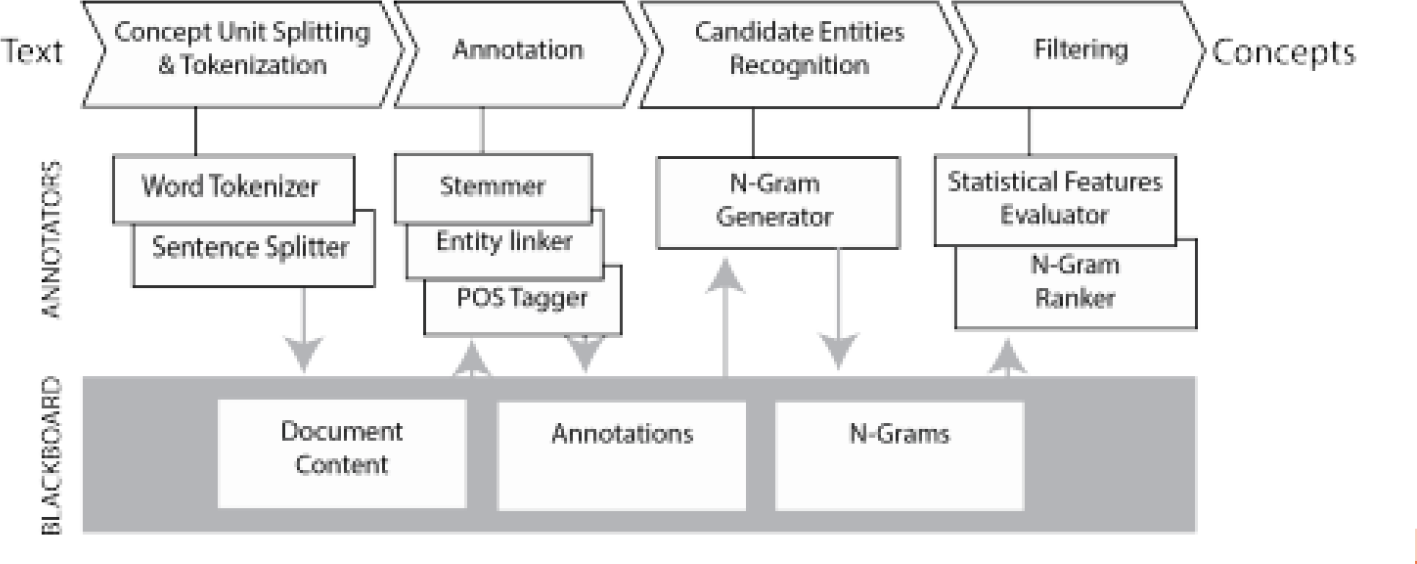
Knowledge extraction pipeline.

To evaluate a deep learning model, as usual, we also used the precision, recall and *F*_1_ *− score* on the test set. By evaluating the set of true positive, false positive and false negative, we computed the value of *F*_1_ *− score*.

## 4 Results and discussion

### 4.1 Result

We assessed the performance of NER tagger method by evaluating the LSTMCRF, on the Oryzabase dataset covering severals different types of rice genomes. LSTM-CRF used as features only low-dimensional representations of the words in the vicinity of the to-be-classified tokens, created by mixing word-level embeddings created in an unsupervised fashion with character-level embeddings trained on the respective corpus. Besides, we also mentioned to the result of the first part of hybrid method which is the dictionary lookup process. Based on this output, we applied some techniques to improve the result.

Results were compared between an original LSTM model; an LSTM with CRF layers which have the same function to recognize entity in the text data using tag information; OGER - a dictionary lookup method which are first part of second method, OGER combining with CRF model ; and OGER with NN.

We evaluated the performance of all models on Oryzabase dataset. Results in terms of precision, recall, *F*_1_ *− score* for each model are shown in table 4. LSTM-CRF achieved the best performance between several models. On average, *F*_1_-score is 86.72% for the generic LSTM-CRF method, 80.44% for the generic LSTM method.

**Table 4.** Result of performance values in terms of precision, recall and *F*_1_ *− score* for pure LSTM and LSTM-CRF method with different training parameters: (i)learning rate=0.001, dropout =0.3, (ii) learning rate=0.001, dropout =0.5.

| | Precision(%) | | Recall(%) | | $F_1 - score$(%) | |
| --- | --- | --- | --- | --- | --- | --- |
|  | (i) | (ii) | (i) | (ii) | (i) | (ii) |
| LSTM | 80.16 | 78.06 | 79.16 | 82.97 | 79.66 | 80.44 |
| LSTM-CRF | 87.24 | 87.32 | 84.73 | 86.13 | 85.97 | 86.72 |

On the other side, the result of hybrid method is very exploitable. The performance of OGER with Conditional Random Field got the best result on testing set among 3 approaches which is 85.08% on average. The second rank is OGER combining with Neural Network, which has reached 67% accuracy on average. Although the improvement are not high as we expected but it still has some improvement rather than the only OGER which is around 58.5%. The result is shown in the table 5.

**Table 5.** Result of performance of hybrid method in terms of precision, recall and *F*_1_ *− score*.

| | Precision(%) | Recall(%) | $F_1 - score$(%) |
| --- | --- | --- | --- |
| OGER | 53.03 | 65.23 | 58.50 |
| OG+NN | 63.93 | 71.10 | 67.32 |
| OG+CRF | 88.39 | 82.24 | 85.08 |

### 4.2 Discussion

During the process to complete the method, we had to deal with several issues coming from the format of data as well as the configuration of the models. About the dataset, the whole data was created from abstract and title of over 10.000 scientific articles in Oryzabase reference list. With the aim to identify gene names or symbols in text data, we expect that this information could appear in the abstract and might be in title. Besides, the huge number of data (2.7 millions words) with different kinds of writing styles takes a lot of time for the pre-processing.

During the training phase, we identified some imbalances of each entity types in the dataset, which had some effects at the performance level for the model. The ratio of information between gene names/symbols but the rest is a big problem. After some training times, the first evaluation of model just reached over 65.5% in accuracy which did not satisfy our expectation. The imbalance of data can lead to worse result during the training process, and the traditional oversampling methods seemed not available for text data, which are really meaningful in the connection of words and sentences.

To handle this problem, we based our training on the workflow of data during NER models, that have input at sequences and process input at word level. Thus, we could collect the all sentences which contain the entity tags without effect the meaning of text data. This solution seemed to be effective to improve the performance of the model when it reaches over 86% in accuracy of *F*_1_ *− score*. In the result section, we shown the results of the performance for a specific parameter which is the value of dropout functions. For both LSTM and LSTMCRF, at the beginning, we set the default value of learning rate equal to 0.005 but these models could not converge. After that, the value of learning rate equal to 0.001 has been chosen through several experiments and evaluation. For the value of dropout, we tried with different values of dropout because we wanted to see the performance of model as well as the effect of parameters to the training process. The default value of dropout equal to 0.5 was firstly chosen for the training process. However, according to Lample *et al*.(2016) [11], the value of dropout equal to 0.3 was optimal for most dataset evaluated. Therefore, we wanted to try for both values and the result is not really effective.

In recent years, pattern- and dictionary-based methods have been replaced by new approaches based on machine learning in general and deep learning more specifically. Nowadays, several methods relying the combination of deep neural network with other techniques are used to develop applications which are able to detect entities automatically rather than manually as the traditional ways. It makes a lot of benefits for the research works as well as for human life. For examples, it can help people to extend the databases which can be used for several purposes of human, or can help scientists finds some new entities. However, scientists normally focus more on the biomedical or somethings that related to human genes. The research in Agronomy now is very popular and plays an important role for human nutrition.

## 5 Future work

Our purpose is finding a method to extract the information of genes, proteins in terms of name or symbols from text data to integrate the database of plant genes for several research purposes. Many different experiments, tests and optimization have been implemented during this project. The result of the model is positive for further research and at this moment, it can be considered as a good baseline method for our project.

In the future, we would like to deploy a couple of optimized ideas so as to improve the result of the current work and start to implement the new methodology to not only improve the accuracy but also optimize the time computing. The following ideas could be tested.

For optimizing the accuracy and time-computing, during the process of building our own deep neural network model to extract biological entities from raw texts that include thousands of scientific paper, we want to apply high performance computing (HPC) on GPU to optimize the computing time of training process for a deep sequential model to handle a large dataset. There would be a challenge when applying parallel computing for the sequential model. However, it is a useful and exploitable direction for future development.

## Footnotes

4 https://shigen.nig.ac.jp/rice/oryzabase/

